## Supplementary material for "Between-day reliability of local and global muscle-tendon unit assessments in female athletes whilst controlling for menstrual cycle phase": S1 and S2 Tables

S1 Table. Definitions and calculations of local muscle-tendon unit metrics.

| **Variable** | **Definition/calculation** |
| --- | --- |
| **Achilles’ tendon metrics** |  |
| Raw elongation (mm) | Maximum elongation of the AT measured from rest to MVC. |
| Passive elongation (mm) | The elongation of the AT due to joint rotation that occurs during maximal contractions. |
| Corrected elongation (mm) | Elongation of the AT from rest to MVC, corrected for the elongation due to passive joint rotation (i.e., raw elongation minus passive elongation). This variable is used for all strain and stiffness calculations. |
| Strain (%) | Elongation of the AT relative to resting length. |
| AT force (N) | Ankle plantar flexion moment (recorded by the dynamometer) divided by the AT moment arm. |
| AT*k* (N/m) | Stiffness – slope of the AT force-elongation curve from 0-100% (AT*k*_all_), 0-50% (AT*k*_low_), or 50-100% (AT*k*_high_) of MVC. |
| NAT*k* (N/strain) | Normalised stiffness – slope of the AT force-strain curve from 0-100% (NA*k*_all_), 0-50% (NAT*k*_low_), or 50-100% (NAT*k*_high_) of MVC. |
| AT*k*_index_ (N/strain) | Maximum tendon force divided by maximum tendon strain [19]. |
| **Single-joint isometric strength** |  |
| Ankle plantar flexion moment (N.m) | Average maximum plantar flexion moment recorded by the dynamometer across all maximal contractions. |
| Knee extension moment (N.m) | Average maximum knee extension moment recorded by the dynamometer across all maximal contractions. |
| AT, Achilles’ tendon; MVC, maximum voluntary contraction. | |

S2 Table. Definitions and calculations of global muscle-tendon unit metrics.

| **Variable** | **Definition/calculation** |
| --- | --- |
| **Countermovement jump metrics** | |
| Jump height from take-off velocity (cm) | *d = v_f_^2^ / 2g*  Where *d* = jump height, *V_f_* = take-off velocity, and *g* = acceleration due to gravity. **Also used for squat jump.** |
| Jump height from flight time (cm) | *d = v_i_t + ½gt^2^*  Where *d* = jump height, *V_i_* = initial velocity, and *g* = acceleration due to gravity, and *t* = half of flight time. **Also used for squat jump and drop jump.** |
| RPD (W/s) | Slope of the power-time curve from the start of the concentric phase (first instant of positive velocity of the centre of mass) to the instant of peak power [43]. |
| RFD (N/s) | Slope of the force-time curve from the start of the braking phase (instant of maximum negative velocity of the centre of mass) to the instant of peak force [27]. |
| Positive impulse (N.s) | Integration of the vertical force-time curve from the instant that force rises above body weight (i.e., start of the braking phase) until take-off [30]. |
| Countermovement depth (cm) | Negative displacement of the centre of mass calculated via double integration (using trapezoid rule) of acceleration time data, which in turn was calculated by dividing force by body mass. |
| Vertical stiffness (N/m) | Peak force divided by countermovement depth [72]. |
| RSImod | Jump height (via flight time method) divided by time to take-off (defined as the time from onset to take-off. [73]). |
| **Squat jump metrics** | |
| RPD (W/s) | Slope of the power-time curve from onset to the instant of peak power. |
| RFD (N/s) | Slope of the force-time curve from onset to the instant of peak force. |
| RFD time bands (N/s) | Slope of the force-time curve from onset to the specified time point (i.e., 50, 100, or 250 ms). |
| Eccentric Utilisation Ratio | CMJ jump height / SJ jump height (via take-off velocity method [61]). |
| **Drop jump metrics** | |
| RSI | Jump height divided by ground contact time [74]. |
| **Isometric midthigh pull metrics** | |
| Impulse time bands (N.s) | Force curve integrated over specific time band (i.e., 50, 100 or 250 ms). |
| RFD time bands (N/s) | Slope of the force-time curve from onset to the specified time point (i.e., 50, 100, or 250 ms). |
| Dynamic Strength Index | CMJ peak force divided by IMTP peak force [62]. |
| RPD, rate of power development; RFD, rate of force development; RSImod, reactive strength index modified; RSI, reactive strength index. | |
